# AraRoot - A Comprehensive Genome-Scale Metabolic Model for the Arabidopsis Root System

**DOI:** 10.1101/2024.07.28.605515

**Authors:** Lohani Esterhuizen, Nicholas Ampimah, Marna D Yandeau-Nelson, Basil J Nikolau, Erin E Sparks, Rajib Saha

## Abstract

Being the first plant to have its genome sequenced, *Arabidopsis thaliana* (Arabidopsis) is a well-established genetic model plant system. Studies on Arabidopsis have provided major insights into plants’ physiological and biochemical nature. Methods that allow us to computationally study the metabolism of organisms include the use of genome-scale metabolic models (GEMs). Despite its popularity, currently no GEM maps the metabolic activity in the roots of Arabidopsis, which is the organ that faces and responds to stress conditions in the soil. We’ve developed a comprehensive GEM of the Arabidopsis root system - AraRoot. The final model includes 2,682 reactions, 2,748 metabolites, and 1,310 genes. Analyzing the metabolic pathways in the model identified 158 possible bottleneck genes that impact biomass production, most of which were found to be related to phosphorous-containing- and energy-related pathways. Further insights into tissue-specific metabolic reprogramming conclude that the cortex layer in the roots is likely responsible for root growth under prolonged exposure to high salt conditions, while the endodermis and epidermis are responsible for producing metabolites responsible for increased cell wall biosynthesis. The epidermis was found to have a very poor ability to regulate its metabolism during exposure to high salt concentrations. Overall, AraRoot is the first GEM that accurately captures the comprehensive biomass formation and stress responses of the tissues in the Arabidopsis root system.

## 1 Introduction

It is well-established that *Arabidopsis thaliana* (Arabidopsis) is a popular model organism in plant biology (Brady et al., 2007; Chen et al., 2004; Smolko et al., 2021; The Arabidopsis Genome Initiative, 2000; W. Xu et al., 2013). Although not economically important, its simple metabolism, short lifespan, and minimal nutrient requirements has made it the ideal model organism to study plant molecular, cellular, and developmental mechanisms (Poolman et al., 2009; The Arabidopsis Genome Initiative, 2000). The sequencing of the Arabidopsis genome provided much-needed insights into plant metabolism, with recent studies focusing on tissue-specific metabolic pathways. When exposing Arabidopsis to altered growth media and stress conditions, the root system acts as the organ to detect and respond to changes in the soil, such as water scarcity and nutrient depletion (Kellermeier et al., 2014; Ngo & Nakamura, 2022; Smolko et al., 2021; H. Xu et al., 2022). The main responsibility of the roots is to take up all required nutrients and water from the soil while also acting as an anchor and providing some storage for excess nutrients (Chowdhury et al., 2022; Smolko et al., 2021). The absence of complex processes such as photosynthesis and respiration in the roots coupled with a simple cellular organization make this an excellent organ for metabolic modeling (Brady et al., 2007; Kellermeier et al., 2014; W. Xu et al., 2013).

As novel plant biology discoveries have continued to be made, scientists looked to the power of machines for help, which brought about the rise of the genome-scale metabolic model (GEM) (C. G. de O. Dal’Molin et al., 2010; de Oliveira Dal’Molin et al., 2015; Poolman et al., 2009). A GEM mathematically formulates the metabolism of an organism where a matrix represents the stoichiometry between the metabolites and reactions involved in the known biological processes as mapped through genome studies, with associated genes mapped through gene-protein-reaction (GPR) relations (Edwards & Palsson, 1999; Mahadevan et al., 2002). A GEM will always have fewer metabolites than reactions since the system is always assumed to operate at steady state. This would make it an underdetermined system, which would either have no solution, or an infinite number of solutions (Edwards & Palsson, 1999; Poolman et al., 2009). A method to reduce the solution space of the model is to introduce ‘omics’ data to the model to restrict the bounds of the reaction fluxes in the model (Gudmundsson & Thiele, 2010; Sinha et al., 2021).

Experimentally obtained omics data such as transcriptome and proteome data can be incorporated into the model through the GPR by using any one of several well-established methods. Some well-known examples include GIMME (Becker & Palsson, 2008), iMAT (Zur et al., 2010), and E-Flux (Colijn et al., 2009). The expression distributed reaction flux measurement (EXTREAM) algorithm is built on the E-Flux method to account for gene abundance in the organism (Chowdhury et al., 2023). A study conducted by Machado and Herrgård (2014) compared 18 different published methods that incorporate transcriptomics to several different GEMs and concluded that in the case of a biomass objective function, E-Flux is the only method that does not replace the biomass objective with an objective that relies on the specific set of gene expression data (Machado & Herrgård, 2014) and EXTREAM only has a significant impact on metabolic predictions in the case of complex gene relations.

Though experimentally determined biomass composition information yields the most accurate growth predictions by a GEM, it is expensive and time-consuming to conduct these experiments, which is why GEMs often assume a biomass composition established for other similar organisms (Chowdhury et al., 2022; C. G. O. Dal’Molin, Brin 2020). The maize root model published by Chowdhury *et al*. (2022) included a biomass reaction that captured the stoichiometry of each metabolite that contributed to biomass formation based on experimental evidence obtained specifically for the maize root system (Chowdhury et al., 2022). The tissue-specific model published by Schroeder & Saha (2020) contained an Arabidopsis root model consisting of a basic biomass composition that including only five metabolites, based on the composition of switchgrass (*Panicum virgatum*) (Schroeder & Saha, 2020b).

Although tissue-specific models of Arabidopsis have been developed regarding whole-plant metabolism, there are currently no tissue-specific GEMs for the Arabidopsis root system that accurately capture biomass composition and secondary metabolism. This work aims to develop a comprehensive GEM for the Arabidopsis root system – AraRoot. Figure 1 summarizes the methodology followed to reconstruct AraRoot, the first GEM capable of capturing the biomass formation and stress response of the Arabidopsis root system. A basic local alignment search tool for proteins (BLASTp) (Altschul et al., 1990) confirmed high gene homology between Arabidopsis and maize (Provart et al., 2016; Saha et al., 2011; Woodhouse et al., 2021), motivating the use of a published maize root model from Chowdhury *et al*. as a blueprint. In addition, AraRoot includes data from established databases such as KEGG (Kanehisa et al., 2014), TAIR (Berardini et al., 2015), and MaizeGDB (Woodhouse et al., 2021). The model was updated to include an accurate biomass reaction and secondary metabolites such as dipeptides, coumarins, and lignols, which include coniferin, esculetin, scopoletin, and scopolin. Because multiple sources were consulted for the biomass composition, it was crucial to understand the effect that changes to the biomass composition and stoichiometry would have on AraRoot. A previously published algorithm was utilized to quantify the sensitivity of the biomass to perturbations in the biomass metabolite coefficients (Dinh et al., 2022). Shadow price values of the biomass metabolites were next calculated to provide further insights into the dependence of the model on metabolite concentrations (Maranas & Zomorrodi, 2016; Schroeder et al., 2020). Results from the sensitivity analysis and shadow price values indicate that sugar components act as overflow metabolites (positive shadow price values) for biomass formation, while cellulose and xyloglucan concentrations have the highest impact on biomass synthesis. Blocked reactions in the model were identified by applying flux variability analysis (FVA). To distinguish between relevant blocked reactions and irrelevant ones that could be removed from the model, blocked reactions that have high homology from the BLAST results remained in the model. An in-house tool known as OptRecon was used to remove thermodynamically infeasible cycles (TICs) and to add missing pertinent reactions from a custom database, while preventing formation of new TICs. Root tissue transcriptomic data was integrated using the E-Flux algorithm (Li et al., 2016) to reduce the solution space and contextualize the root model. Our recently developed metabolic bottleneck analysis (MBA) was applied to the model to identify the bottleneck reactions (and the associated bottleneck genes) that have an impact on biomass production when strict bounds are imposed to the reaction fluxes (Chowdhury et al., 2023). The 158 potential bottleneck genes identified were further analyzed to gain insights into the correlations that exist between these genes when Arabidopsis roots are grown in normal and high salt conditions.

**Figure 1:**
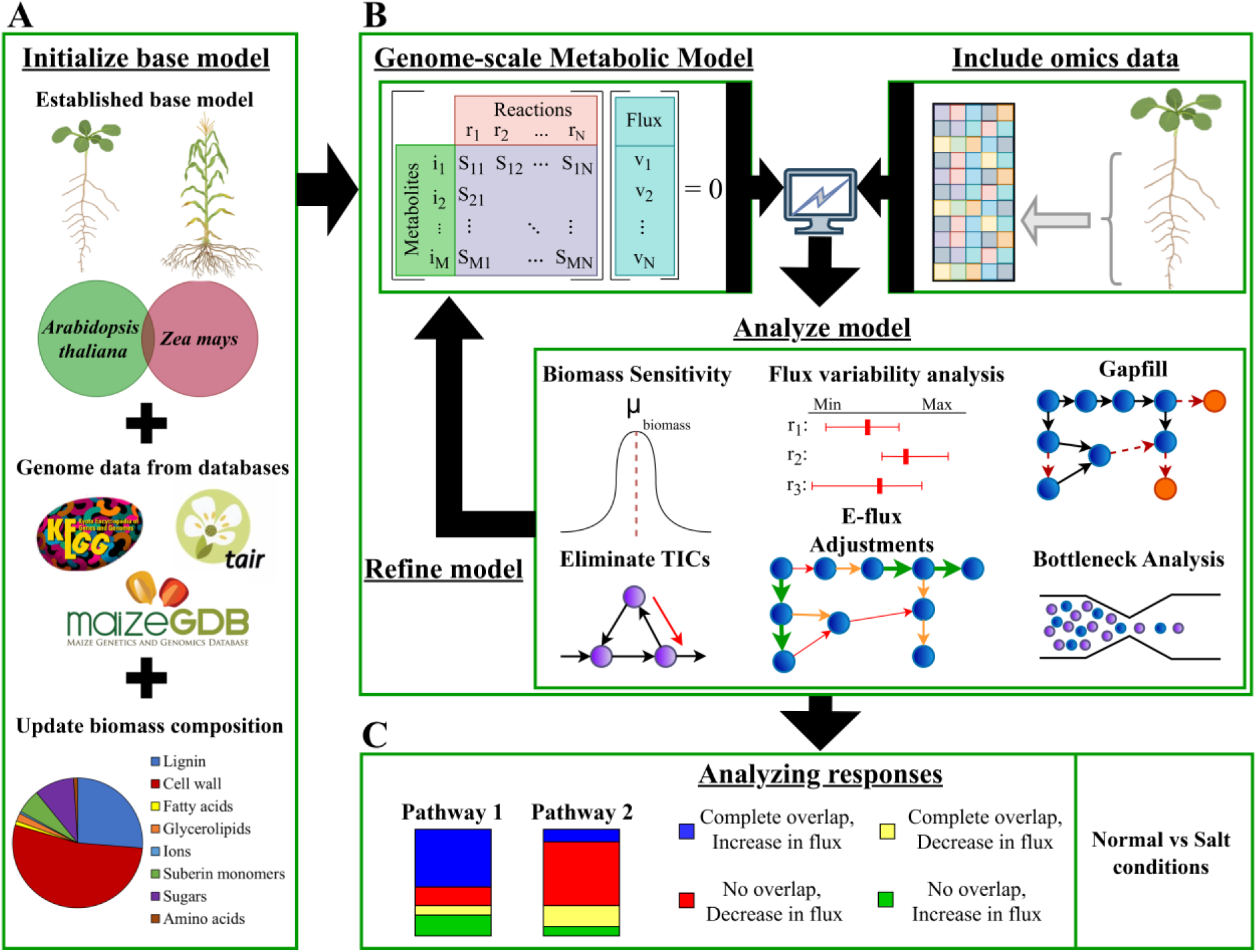
Reconstructing and analyzing the AraRoot model by (A) updating a base root model (*Zea mays*) to include a comprehensive Arabidopsis root biomass and reactions from well-known databases, (B) constructing the genome-scale metabolic model (GEM) and including published transcriptomics to the model, which was analyzed using established methods, and (C) analyzing the response of the model to stress-induced environmental conditions by comparing the flux range variations under normal and salt stress conditions.

Once the AraRoot model was built, it was used to explore the predictive capabilities of the model under root stress responses. Published gene expression data (Geng et al., 2013) was utilized to create cell-type-specific models of the cortex, endodermis, epidermis, and stele tissues under normal and stress conditions using the AraRoot model as a blueprint. Subsequently, the same transcriptomics data were used to analyze the bottleneck genes, thus identifying correlations between the bottleneck genes under stress conditions to limit biomass production. A flux range variation analysis of each cell-type under normal and high salt growth conditions captured the metabolic reprogramming of each tissue during stress-induced conditions. Resulting variations in reaction flux ranges indicate that the cortex ultimately drives growth recovery in the roots after prolonged exposure to salt, with increased reaction flux ranges found in most of the metabolic pathways. The epidermis and endodermis were found to be responsible for producing more metabolites related to cell wall biosynthesis with flux ranges in the pentose phosphate pathway increasing during salt exposure, particularly the production of D-ribulose 5-phosphate. Results also indicate that the epidermis lacks the ability to further regulate metabolism during stress-induced growth. Together, this work proposes a tissue-specific GEM of the roots, capable of capturing metabolic variation under different growth conditions.

## 2 Methods

### 2.1 Initializing the model

The reconstruction and analysis of the AraRoot model aims to produce a GEM capable of accurately capturing the biomass formation and stress response of the Arabidopsis root system. Model initialization involves establishing a base model capable of reconstruction to represent an Arabidopsis root system. Previous literature indicates genetic similarity between maize and Arabidopsis in terms of metabolic pathways (Chowdhury et al., 2022; Saha et al., 2011). The protein sequences of genes in the Arabidopsis root system, as established by Li *et al*. (2016), were obtained from the TAIR database and were set as the query sequences for BLASTp. The protein sequences of the genes included in the published maize root base model were obtained from the KEGG database and were considered the subject sequences. The screening criteria for the BLASTp result hits were set at a percentage identity (PID) of 60%, a query coverage (COV) of 0.8, and a Max Score of 400.

The biomass composition of the Arabidopsis root system was determined by consulting literature. A complete list of all metabolites and their respective compositions that make up the biomass can be found in Supplemental File 1, accompanied by links to the literature sources. The biomass reaction in the base model was updated according to the new biomass, and several well-known techniques were used to ensure that the base model could produce biomass with the updated composition. These techniques include flux balance analysis (FBA) and flux variability analysis (FVA). The mathematical formulation of the optimization problem used in both techniques can be found in Supplemental File 2.

Once the biomass reaction was updated, a previously published algorithm was used to analyze the sensitivity of the model to perturbations in the biomass metabolite coefficients (Dinh et al., 2022). The proposed algorithm calculates a standard deviation ratio (SDR), which quantitively determines the impact that deviations in the input parameter *c* have on the output fluxes *ν* of a GEM using the following equation:

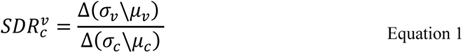

Where *σ* and *µ* represents the standard deviation and mean, respectively. Parameters were sampled 10,000 times from a normal distribution with relative standard deviations of 5%, 10, and 20% centered around the original values. Noise was imposed on all coefficients simultaneously, followed by imposition on cellulose and xylan singularly as biomass metabolites. For further insights into the relationship between the metabolites and biomass formation in the base model, the shadow price values (*λ*) of the metabolites were calculated by considering the dual problem of the FBA formulation. An upper and lower shadow price range was determined by utilizing the coefficients of the biomass reactions determined at 20 % perturbations during the biomass sensitivity analysis. Negative shadow price values would indicate growth-limiting metabolites, which are metabolites whose availability indirectly affect biomass formation. Positive shadow price values are associated with overflow metabolites, which are metabolites whose availability directly affects biomass formation. The mathematical formulation for calculating the shadow price value of each metabolite can be found in Supplemental File 2.

### 2.2 Model reconstruction

After incorporating the updated biomass reaction to the base GEM, a standard model reconstruction procedure was followed with added steps to capture comprehensive root metabolism (Thiele & Palsson, 2010). It was necessary to determine which reactions in the base model had become redundant given the new biomass reaction, since some reactions in the model were only specific to maize and would also need to be removed. FVA was utilized to identify all blocked reactions, which are reactions that did not carry flux during biomass production. The blocked reactions were compared to reactions catalyzed by specific gene products that showed high sequence homology between Arabidopsis and maize. Common reactions remained in the model, while blocked reactions with no homology were removed from the model.

An in-house tool known as OptRecon was utilized to identify and remove thermodynamically infeasible cycles (TICs) (Nelson & Saha, 2024). The mathematical formulation capable of identifying TICs in a GEM is provided in Supplemental File 2 and forms the basis for OptRecon. To account for missing reactions from the maize base model, a custom database of reactions was created from which relevant reactions were added. The custom database was created by comparing the transcriptomes of the Arabidopsis root system to the published root system transcriptomes of other similar C3 plants, including tomato (Pirona et al., 2023), soybean (Adhikari et al., 2019), and rice (Liu et al., 2021). Reactions that were found to have high homology between Arabidopsis and the three other plants were compiled to form a custom database, and OptRecon was utilized to incorporate the maximum number of reactions without creating any TICs.

To reduce the solution space and contextualize the model, gene expression data was incorporated into the model. The gene-protein-reaction (GPR) relation of Arabidopsis as published by Saha *et al* (2011) was consulted to include transcriptomics data from the maturation zone of the roots obtained by Li *et al*. by applying the E-Flux algorithm (Sinha et al., 2021). Metabolic bottleneck analysis (MBA) identified bottleneck reactions that had formed because of incorporating transcriptomics into the model. Figure 2 provides a flow diagram that shows how MBA is performed on a GEM to identify bottleneck reactions by expanding flux bounds and determining new maximum biomass flux values using FBA. The new optimal value (NB) is compared to the initial maximum biomass (B), and if the solution changes, the reaction *j* is considered a bottleneck reaction.

**Figure 2:**
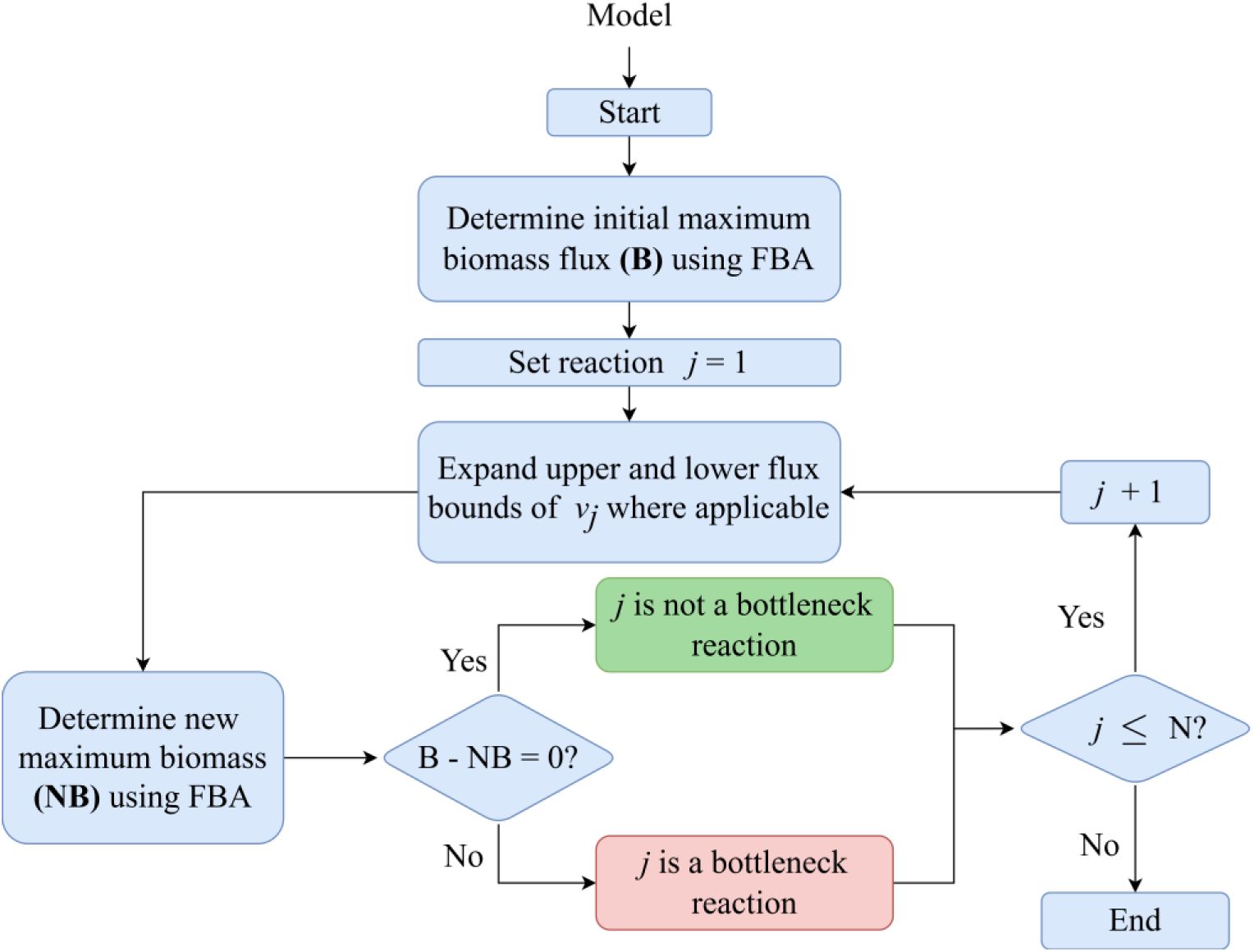
Flow diagram for performing metabolic bottleneck analysis (MBA) to identify bottleneck reactions in a GEM.

The associated bottleneck genes were identified from the same Arabidopsis GPR, and gene expression values from published results were used to gain further insights into the nature of the bottleneck genes. The transcriptomics data were collected for the cortex, endodermis, epidermis and stele root tissue under normal growth conditions and stress-induced growth conditions (Geng et al., 2013). The bottleneck genes and gene expression values of each tissue were compared using the t-distributed Stochastic Neighbor Embedding (t-SNE) clustering analysis to evaluate transcriptomic correlations between bottleneck genes under normal growth conditions and stress-induced growth conditions. The genes associated with each cluster in the t-SNE analysis was further studied using protein-protein interaction networks available in the STRING database (Szklarczyk et al., 2023).

### 2.3 Biological insights

Dynamic transcriptomic datasets were used to analyze flux range variability of the Arabidopsis seedlings (Columbia-0 accession) exposed to media supplemented with 140 mM of sodium chloride (NaCl; salt) (Geng et al., 2013). Spatiotemporal micro-array-based transcriptional profiles were determined for the cortex, endodermis, epidermis, and stele tissues of the root system. The gene expression data collected from the different tissues in normal growth conditions, 1 hour of salt exposure, and 48 hours of salt exposure were incorporated into AraRoot using the E-Flux method to create tissue-specific root models for each of the growth conditions. The normal growth condition was considered as a baseline for the flux ranges while the flux ranges resulting from the 1-hour exposure and 48-hour exposure were compared to the baseline to compare the variations in flux ranges during high salt concentration growth conditions. Figure 3 lists the categories of flux variation changes that can occur under high salt concentration growth conditions. The flux ranges found under 1-hour salt exposure were first compared to the flux ranges found under normal conditions, followed by also comparing the 48-hour salt exposure to the normal conditions.

**Figure 3:**
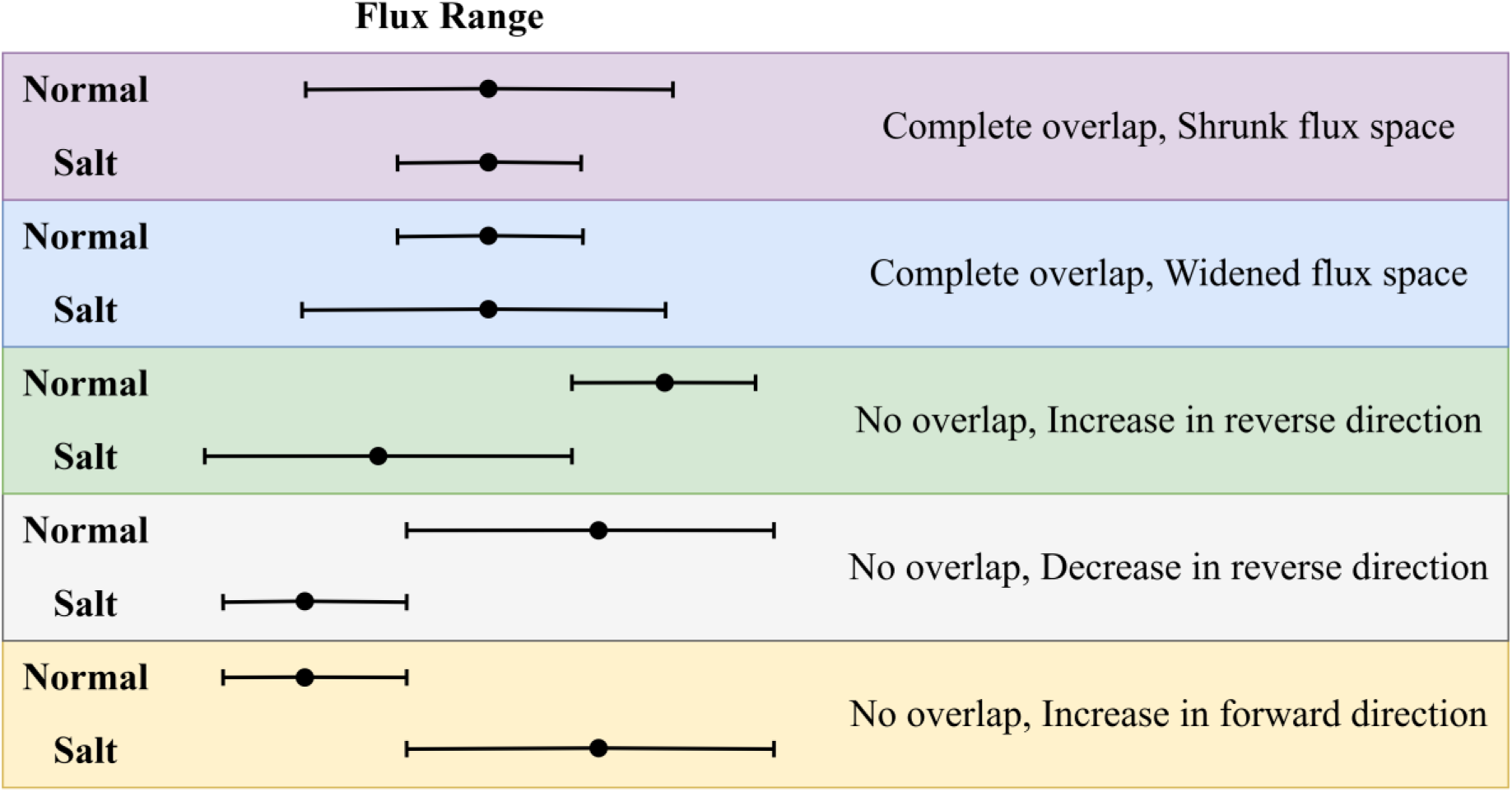
Five categories to classify results of flux variation analysis. Reaction flux ranges of the stress condition (Salt) are compared to the base condition (Normal).

Shrunk flux spaces under high salt concentration conditions indicate pathways that produce fewer metabolites under reduced flux, while widened flux spaces indicate an increase in metabolite production. These conditions are a direct result of the reactions associated with under-expressed and over-expressed genes respectively. Variations in flux range with no overlap indicate major changes in metabolic reprogramming under stressful growth conditions. The variations in flux ranges can be confirmed using previously published findings in changes in metabolite production under high salt growth conditions.

### 2.4 Simulation software

All simulations were run on a combination of General Algebraic Modeling System (GAMS) version 24.7.4 and Python version 3.10. In both cases the IBM CPLEX licensed solver was used for optimization calculations such as FBA, MBA, and FVA. All simulations were run on a Linux-based terminal connected to the Holland Computing Center at the University of Nebraska to allow for decreased computational time and adequate memory to run extensive optimization formulations. The final version of AraRoot is available on the SSBio GitHub page (https://github.com/ssbio).

## 3 Results

### 3.1 Model initialization

This work presents AraRoot, the first GEM that aims to accurately capture the biomass formation and stress response of the Arabidopsis root system. The model consists of 2,682 reactions, 2,748 metabolites, and 1,310 genes and was analyzed to determine the metabolic reprogramming that occurs in the root tissues during exposure to stress. Considering Figure 1, the first step was initializing the model, which involves cross-referencing well-known databases for information on the genome, as well as consulting previously published models of a similar nature. The maize root model published by Chowdhury *et al*. (2022) was considered a good candidate base root model given the major similarities that exist between the protein sequences obtained from maize and Arabidopsis (Dukowic-Schulze et al., 2014; Saha et al., 2011; Schroeder & Saha, 2020a).

To verify the feasibility of using the maize root model as a base model, the genes from the maize and Arabidopsis root systems and their associated protein sequences were included in a BLASTp analysis. Results showed 960 hits between the protein sequences of the two root systems according to the cut-off criteria set forth in Section 2.1, indicating that the metabolic system is likely to be highly similar between both root systems. This justifies the use of the previously published maize root model as a base model for AraRoot. Using the maize root model as a base model, the first step involved updating the composition and stoichiometry of the biomass reaction to accurately reflect the biomass of the Arabidopsis root system rather than that of the maize root system. The most noticeable modification to biomass composition in this model includes the addition of cell wall components (cellulose, hemicellulose, and pectin) and the increase in lignin composition, both of which are responsible for structural support and strength in the roots. The specificity of including suberin monomers in the biomass instead of lipids has been well established in several root studies (Pirona et al., 2023; Soltis & Soltis, 2021). The mass composition of the lignin, pectin, cellulose, and hemicellulose was obtained from the C3 root composition analyses (Saunders et al., 2006). The specific split between p-hydroxyphenyl lignin, guaiacyl lignin, and syringyl lignin was assumed to be like that of maize.

Hemicellulose as a compound is a complex branched polysaccharide that can vary significantly between plant species (Rao et al., 2023; Zabotina et al., 2012). A study by Zabotina *et al*. (2012) shows that the hemicellulose in Arabidopsis mainly consists of xylan and xyloglucan, with xyloglucan making up the majority of the hemicellulose composition. Since very little is known about the reaction mechanism of hemicellulose formation, hemicellulose is represented by xylan and xyloglucan in the root model. Saunders *et al*. (2006) was able to quantify the mass composition of the total sugars, fatty acids, amino acids, and suberin monomers found in the root systems of a collection of mostly C3 plants. The sugar compounds and compositions were assumed to be similar to the root system of Arabidopsis proposed by Schroeder & Saha (2020). The specific weight percentage composition of each fatty acid and suberin monomer was determined by previous experimental studies (Li-Beisson et al., 2013). The specific g/gDW compositions of the amino acids in the roots were assumed to be similar to that in the root system of tomatoes, another C3 class plant (Gerlin et al., 2022). Figure 4A illustrates the component groups that are present in the biomass composition of the root system, and all biomass components and their associated mass percentage compositions can be found in Supplemental File 1. The cell wall components, pectin and lignin make up most of the root biomass, while the ions involved in biomass formation were reduced compared to the maize root biomass.

**Figure 4:**
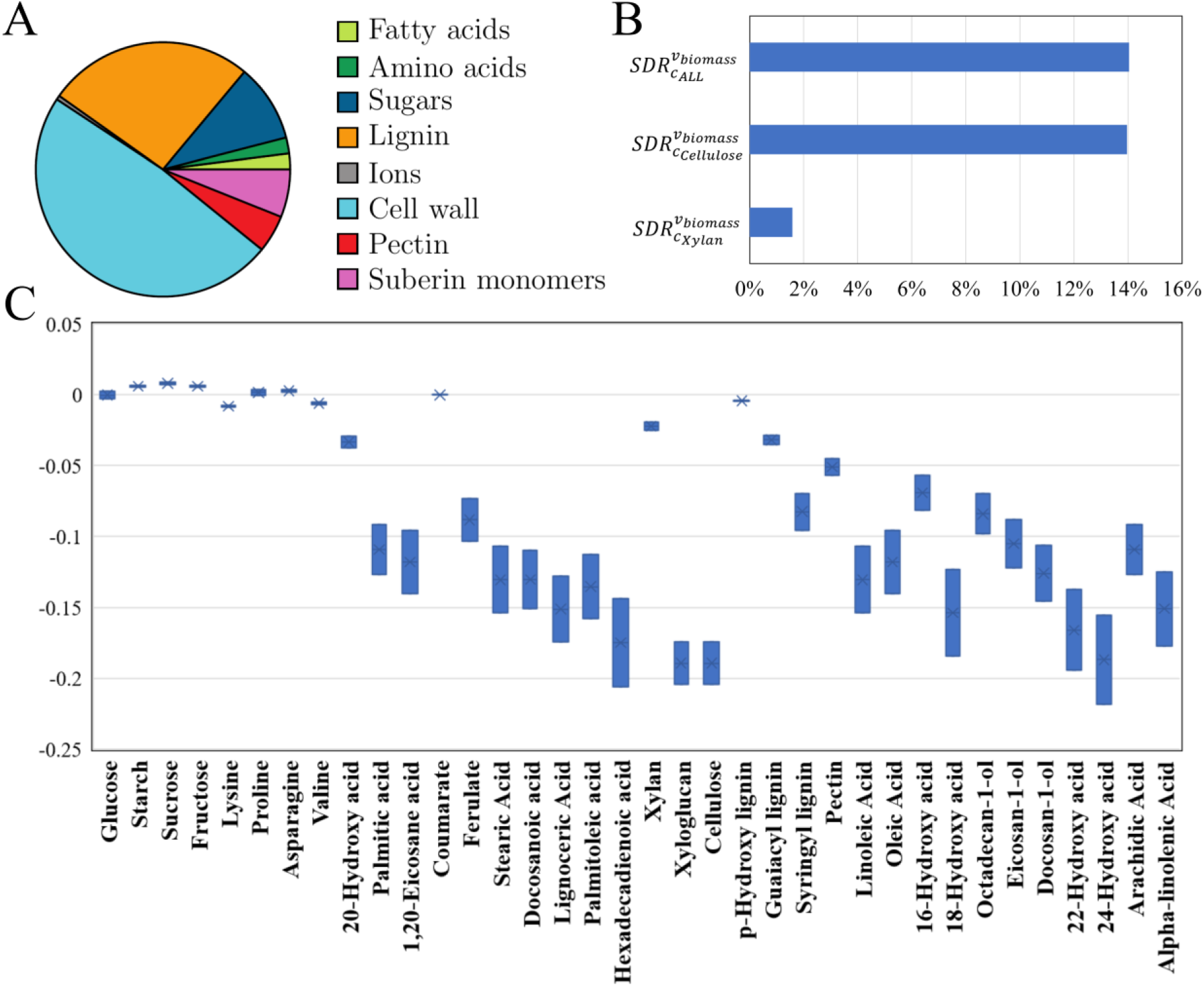
Update and analysis of the Arabidopsis root biomass composition, including (A) the major component groups included in the model, (B) the results of the quantified uncertainty in changes to the biomass coefficients in the model, and (C) the range of shadow price values for the biomass metabolites when coefficient values in the biomass reaction are deviated by 20%.

Once the biomass reaction in the initial model was updated by changing the stoichiometry and components to match the biomass composition, some reactions had to be added for the model to be able to carry flux through the biomass reaction. The reactions needed and their associated mechanisms were found using the KEGG database (Kanehisa et al., 2014). Since all biomass components and their compositions were obtained from literature, a sensitivity analysis was conducted to quantify the uncertainty of the coefficients of the biomass components in the base model by implementing the algorithm developed by Dinh *et al* (2022). Figure 4B provides the standard deviation ratio (SDR) results of the biomass coefficient sensitivity analysis where fluctuations in coefficient values were imposed to all coefficients at once, only on cellulose, and xylan as singular coefficients. The temporal fluctuations imposed on all coefficients at the same time (*c*_*ALL*_) shows a significant impact on the predicted biomass flux through FBA at an SDR of 14.04%. Imposing a fluctuation on the coefficient of cellulose (*c_Cellulose_*), a major component in the biomass, also results in a significant impact on the biomass predictions with an SDR of 13.96%. The results for imposing fluctuations on the xylan coefficient (*c*_*Xylan*_) shows little impact on the biomass formation with an SDR of 1.58%. Expanding on the biomass coefficient sensitivity analysis, the shadow price values of all metabolites in the model with the updated biomass reaction was calculated. To determine the range of shadow price values for the metabolites, the coefficients of the biomass reaction were changed to reflect a 20% fluctuation in the coefficient values. Figure 4C provides the shadow price value ranges for all the biomass metabolites.

The positive shadow price values of the sugar components suggest that they act as overflow metabolites, as do proline and asparagine. All other biomass metabolites were found to have negative shadow price values, indicating that they act as growth-limiting metabolites. The shadow price values calculated for hexadecadienoic acid, xyloglucan, cellulose and 22-hydroxydocosanoic acid indicate that these metabolites would have the highest correlation to biomass synthesis in the roots. Hexadecadienoic acid (fatty acid) and 22-hydroxydocosanoic acid (suberin) make up a minute part of the biomass composition, while cellulose and xyloglucan within the cell wall make up relatively large portions of the biomass composition.

### 3.2 Model reconstruction

#### 3.2.1 Resolved blocked reactions and TICs

At this stage, the base model included 4,089 reactions, 4,419 metabolites, and 6,389 genes. Conducting FVA on the model identifies blocked reactions by identifying all the reactions that have a maximum and minimum optimal flux of zero when producing biomass in the case of AraRoot. The base model was found to have 2,315 blocked reactions. To determine which of the blocked reactions needed to remain in the model due to possible downstream involvement in pathways, the list of blocked reactions was compared to the reactions that are associated with genes that have high homology between Arabidopsis and maize as obtained from the BLAST results. From the comparison, 678 reactions were found to have high homology between Arabidopsis and maize and would need to remain in the model, despite not carrying flux during biomass production. The remaining 1,637 blocked reactions were then manually scrutinized to ensure that downstream reactions were also kept in the model. From the remaining reactions, an additional 22 reactions were found to play a metabolic role in Arabidopsis and were kept in the model. The remaining blocked reactions were removed from the model.

After the removable reactions had been excluded, all missing secondary metabolites and their associated missing reactions were added to the model. Secondary metabolites that were added include coniferin, syringin, lariciresinol, esculetin, scopoletin, esculin, and scopolin. Once the secondary metabolites and reactions were added to the model it could be evaluated for any TICs. Since a previously published maize root model was used as the base model for AraRoot the inherent base model did not contain any TICs, as confirmed by the tool, OptRecon. Once this was confirmed, it was necessary to consider that the model might be missing some reactions that could be present in Arabidopsis but would be missing from the maize root model.

To find and add any missing reactions, a custom database containing reactions from the whole-plant Arabidopsis genome was created. The SBML file containing all the custom database reactions can be found in the AraRoot GitHub repository (https://github.com/ssbio/AraRoot), and the compilation of the genes used to create the custom database can be found in Supplemental File 4. These reactions were identified by obtaining transcriptomes data of the root systems of other C3 plants, specifically rice, tomato, and soybean. The genes that are expressed in these root systems were found to have homologous genes that are expressed in Arabidopsis roots. Referring to the gene-protein-reaction (GPR) relationship of Arabidopsis, a list of reactions was identified that are associated with genes that have high homology among C3 root systems. These reactions were considered to form the custom database from which OptRecon could add reactions to account for missing reactions. Figure 5 illustrates the steps taken to create the custom database and use OptRecon to fill the missing reactions into the model, constructing the final AraRoot model.

**Figure 5:**
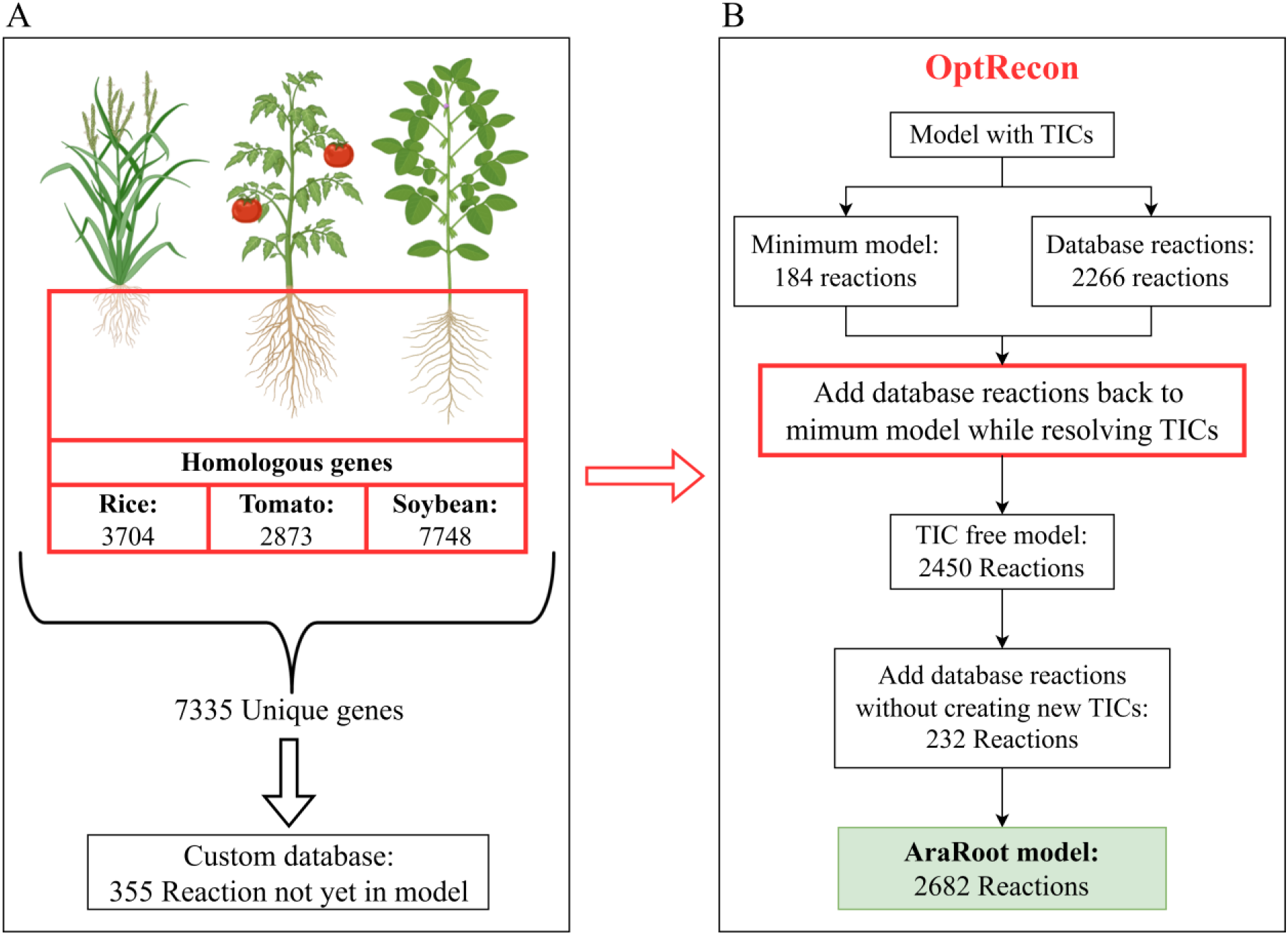
Resolving missing reactions in model by (A) creating a custom database from other C3 plants, followed by (B) using OptRecon to add reactions to the model from a custom database without creating any TICs.

A list of 7,335 unique Arabidopsis genes were found to have high homology with the root systems of rice, tomato, and soybean. From these genes, 355 reactions were identified from the published *A. thaliana* iRS1597 (Saha et al., 2011) model GPR and were used as a custom reaction database for OptRecon. From the custom database, 232 reactions were added to the model. The final AraRoot model consists of 2,682 reactions, 2,748 metabolites, and 1,310 genes.

#### 3.2.2 The inclusion of transcriptomics

At this stage the bounds of the fluxes in the model were all set to large values, essentially “unbounding” the flux ranges. When computing FBA with unbounded flux ranges, the possible solutions to optimizing the maximum biomass flux (*ν*_*biomass*_) are infinite and can take up very large unrealistic fluxes through the reaction pathways in the model. The best practice to reduce the solution space and contextualize the model is to include empirical transcriptomic data of the Arabidopsis root system into the model. The constricted flux bounds will also result in more realistic fluxes through the metabolic pathways as compared to what happens in nature.

Gene expression data obtained from literature was incorporated into the model to contextualize the reaction bounds to more realistic limits. The work by Li *et al*. (2016) generated a high-resolution spatiotemporal map of the gene expressions in the Arabidopsis root system. The fragments per kilobase per million mapped fragments (FPKM) values of 23,934 unique genes were obtained for the meristematic, elongation, and maturation zones of the root system. From the Arabidopsis GPR, 1,310 unique genes are associated with 1,171 reactions from the whole-plant Arabidopsis genome. Of the 1,171 gene-associated reactions, 507 were present in the AraRoot model and the associated bounds were adjusted using the E-Flux algorithm.

#### 3.2.3 Metabolic bottleneck analysis results

Following the procedure for MBA as described in Figure 2, 106 bottleneck reactions were identified, which correlate with 158 unique Arabidopsis genes according to the GPR. Further information on the nature of these genes was gathered to better understand their nature. Transcriptomic data collected by Geng *et al*. (2013) included gene expression values of four different tissues in the root under normal growth conditions and high salt growth conditions. The tissue-specific transcriptomic data was included in the model through E-Flux, and t-Distributed Stochastic Neighbor Embedding (t-SNE) clustering analysis revealed further insights into the correlations in expression between the proposed bottleneck genes in normal and salt conditions. Figure 6 provides the clustering results found for each cell type in the root to determine the metabolic correlations between the bottleneck genes when the Arabidopsis roots are exposed to high salt concentration growth conditions.

**Figure 6:**
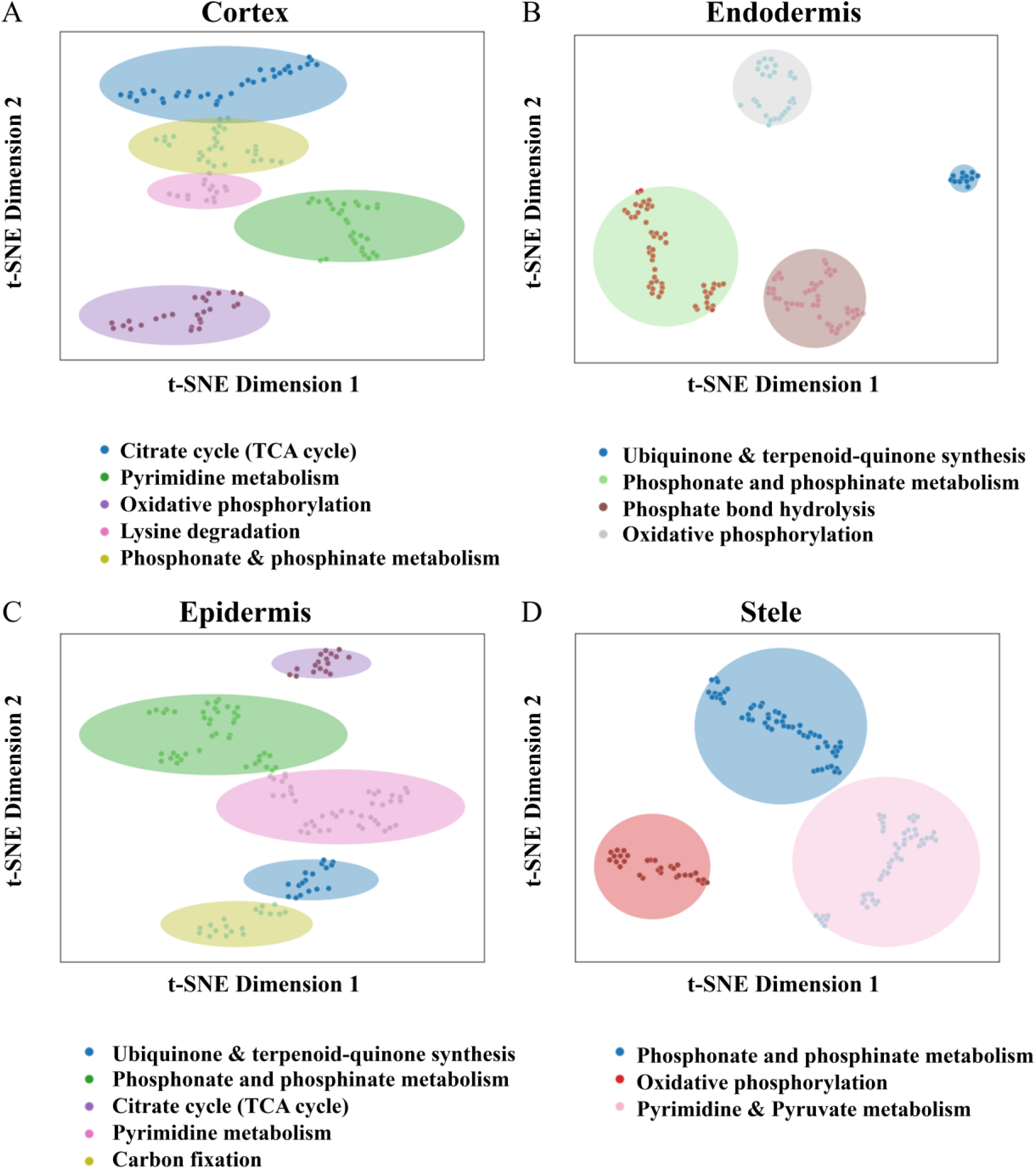
K-Means clustering analysis results of bottleneck genes in the (A) Cortex, (B) Endodermis, (C) Epidermis, and (D) Stele to indicate genetic correlations of the bottleneck genes in normal and salt conditions.

In the endodermis and stele, the bottleneck genes were found to form distinct clusters where the majority of the genes in both cell types are associated with the oxidative phosphorylation, and phosphonate and phosphinate metabolism pathways. Phosphonate and phosphinate metabolism are also prevalent in the cortex and epidermis, though the clusters are not as distinct as in the endodermis and stele. Several bottleneck genes were found to impact the TCA cycle and pyrimidine metabolism in the cortex and epidermis in high salt concentration growth conditions. The bottleneck genes impacting the ubiquinone and other terpenoid-quinone biosynthesis pathways in the endodermis and epidermis formed a distinct cluster in each case, indicating a high correlation during exposure to normal and salt conditions. The cortex seems to be the only cell type with bottleneck genes involved in the lysine degradation pathway, which is a downstream pathway of the TCA cycle. The bottleneck genes and their associated gene expression values in each identified cluster were imported into the STRING database and the protein-protein interaction networks found for Arabidopsis provided further insights into the correlations between the bottleneck genes in each root tissue under the salt stress growth conditions. The illustrative outputs of the protein-protein interaction networks can be found in Supplemental File 3. From comparing the protein-protein networks of each tissue to one another it was found that some proteins found in each of the four networks include PBL1, ALA2, PECT1, and CCT1.

### 3.3 Biological insights

To better understand the capabilities AraRoot has for capturing the metabolic preprogramming that occurs in stress conditions, a flux range variation analysis was carried out to investigate the changes in reaction fluxes in the roots when exposed to high salt concentration growth conditions. The transcriptomic data published by Geng et al. (2013) included gene expression values of the cortex, endodermis, epidermis, and stele tissues in the root system that were exposed to normal growth conditions, 1 hour of salt exposure (short exposure), and 48 hours of salt exposure (long exposure). This data was implemented into AraRoot through the E-Flux algorithm to formulate 12 different tissue-specific models under the various growth conditions. The flux ranges found under normal growth conditions were considered the baseline fluxes, and the flux ranges that resulted from the high salt concentration growth conditions were each compared to the normal growth condition flux ranges according to the categories described in Figure 3. The resulting variations in flux ranges from normal growth conditions to high salt concentration growth conditions is provided in Figure 7.

**Figure 7:**
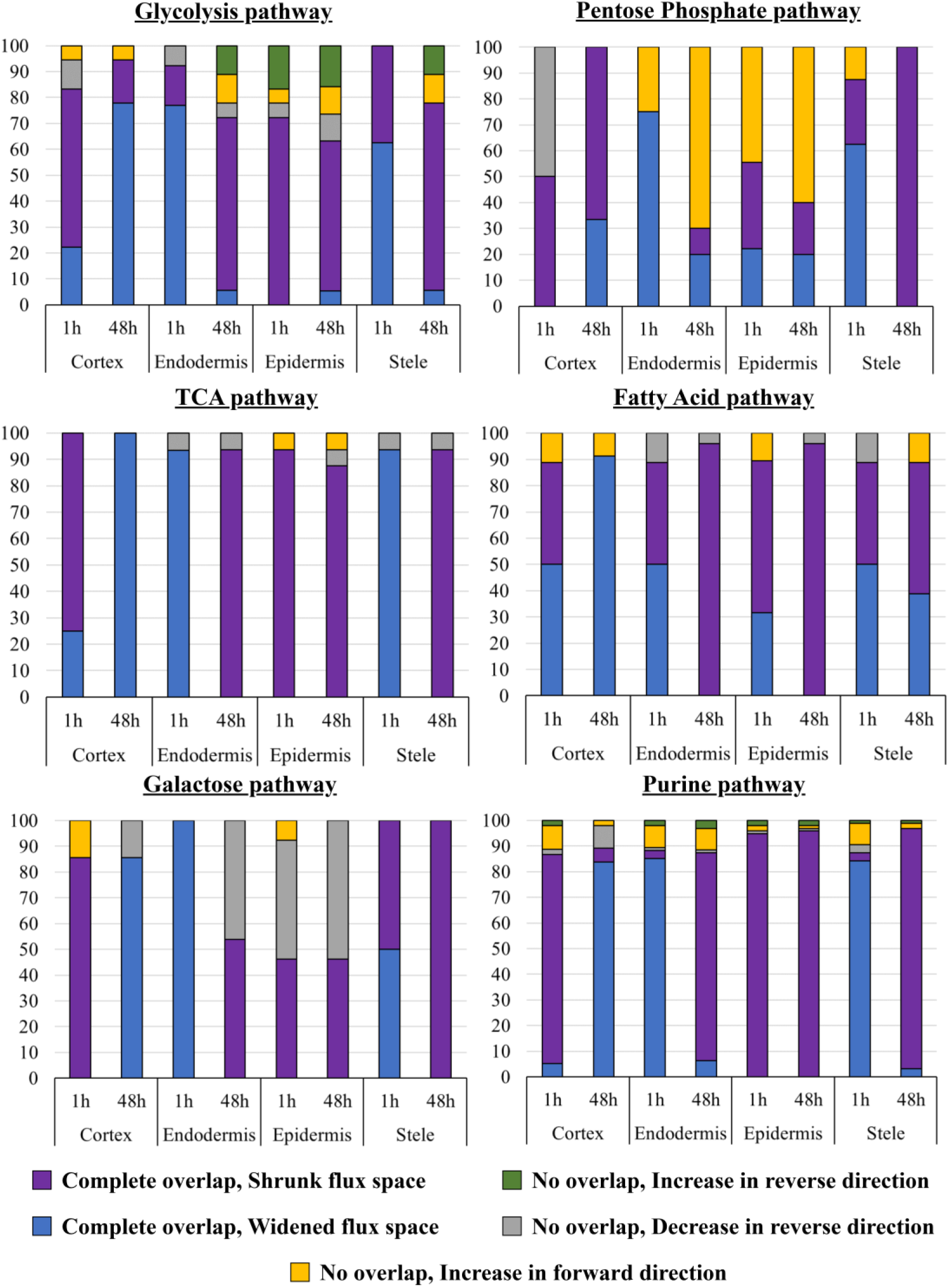
Flux range variation resulting from high salt exposure during root growth.

Central carbon metabolism includes the glycolysis, pentose phosphate, and TCA cycle pathways, which are responsible for the conversion of carbon-containing compounds to sugars and organic molecules and is perhaps the best starting point when analyzing the metabolic reprogramming of the roots under salt stress (Dieuaide-Noubhani et al., 2007; Nägele et al., 2010). Other major pathways that appear upstream and downstream from the central carbon metabolism include the fatty acid, galactose, and purine pathways.

The majority of the pathways in the cortex experience a shrunk flux space after 1 hour of salt exposure, but prolonged salt exposure led to widened flux spaces in all pathways except the pentose phosphate pathway. The endodermis and stele seem to experience the opposite metabolic reprogramming, experiencing widened flux spaces after 1 hour salt exposure while prolonged salt exposure leads to shrunk flux spaces in all pathways except the pentose phosphate pathway. The epidermis seems to experience overall shrunk flux spaces in both short and long salt exposure times.

## 4 Discussion

### 4.1 Reconstructing AraRoot

This work introduces AraRoot, a GEM of the Arabidopsis root system that includes a comprehensive biomass composition that for the first time has the capability of capturing the metabolic reprogramming that occurs in separate root tissues when exposed to stress-induced growth conditions. The development of tissue-specific GEMs such as AraRoot is crucial for gaining insights into the unique metabolic functions and adaptations of distinct plant organs, such as the roots. AraRoot successfully allows for a more precise understanding of how specific tissues, including the cortex, endodermis, epidermis, and stele, respond to environmental stimuli, which advances our knowledge in plant biology and can be further expanded to other crop GEMs to gain insights on how to improve crop resilience.

Our in-house tool, OptRecon, was instrumental in the development of AraRoot. OptRecon facilitated the accurate reconstruction of the Arabidopsis root metabolic network by removing TICs and ensuring completeness of the model by adding missing reactions from a database. This tool enabled us to refine the metabolic pathways specific to the different root tissues, thereby enhancing the predictive power and reliability of AraRoot. The successful application of OptRecon in this study underscores its potential as a valuable resource for developing other tissue-specific GEMs, further contributing to the field of plant systems biology.

Another crucial part of developing AraRoot was ensuring that the model accurately captures a realistic biomass composition of the root organ. An accurate biomass composition would provide a realistic representation of the metabolic demands and outputs of the roots, leading to more accurate simulations and predictions of metabolic behavior under various conditions. The integration of such precise biomass data ensures that the model can better replicate the physiological and biochemical states of tissues under different growth conditions. The biomass composition included in AraRoot was determined from several literature sources and the sensitivity analysis of the model to changes in the biomass composition and stoichiometry highlights the need to experimentally verify the actual biomass composition of Arabidopsis roots. The shadow price values of the biomass components indicate that most of the sugar components act as overflow metabolites. When considering the starch and sucrose metabolism pathways, glucose is formed from D-glucose 1-phosphate, which is also needed to form the cellulose and hemicellulose components of the cell wall. Since cellulose makes up most of the biomass composition, higher flux is required through the cellulose formation reaction, resulting in glucose being an overflow metabolite. The negative shadow price values indicate growth-limiting metabolites. From the shadow price results, cellulose and xyloglucan within the cell wall would impact the actual biomass composition more than any of the other biomass metabolites since they have highly negative shadow price values and make up a relatively large proportion of the biomass composition. The shadow price value of cellulose ties in with the sensitivity analysis of the model after perturbations in the cellulose coefficients, both indicating that changes to the amount of cellulose in the biomass composition could highly affect the model outcomes. The shadow price value of xylan was found to be close to zero, while also indicating that it is a growth-limiting metabolite. This corresponds with the results of the sensitivity analysis with a SDR of 1.58%, both indicating that changes in the xylan composition of the biomass would not have a significant effect on the biomass predictions by AraRoot.

Additionally, the fatty acids, suberin monomers, and lignin components of root biomass all exhibit growth-limiting properties. Both the fatty acids and the suberin monomers depend on acetyl-CoA as their major precursor metabolite and are dependent on flux through the fatty acid and cutin, suberin, and wax biosynthesis pathways for producibility. Although the fatty acids and suberin monomers make up a smaller proportion of the biomass, perturbations in the coefficients of the components affect the other components in the groups as well, creating a "ripple effect" in the biomass production predictions, making these metabolites growth-limiting metabolites. The insights gained from analyzing the bottleneck genes identified from the model provide a unique perspective on the metabolic behavior of the root system and would have taken much longer to gain from empirical investigations, highlighting the incredible power of utilizing tissue-specific predictive GEMs. Developing the GEMs for the root systems of other crops could provide significant insights into the metabolic behaviors of those root systems under stress conditions, allowing for the identification of how to make crops more resilient by overcoming bottlenecks in metabolism.

### 4.2 Predicting stress responses

Developing the AraRoot GEM before contextualizing it to represent specific root cell-types provided a foundational framework that encompasses the comprehensive metabolic capabilities of the root. This includes all known metabolic reactions and pathways within the root, providing a holistic understanding that can be selectively refined and tailored to reflect the metabolic activities of specific cell types. Importantly, the whole-root GEM could also be contextualized to represent any tissue-specific root metabolism. Furthermore, the development of a whole-root GEM allows for the utilization of a wide range of experimental data, including transcriptomics, proteomics, and metabolomics data, and allows for easy adaptation or expansion as new information becomes available in the future. To analyze the capability of AraRoot in capturing the response of the root system to stress, the model was contextualized to represent the cortex, endodermis, epidermis, and stele tissues in the root system. Publicly available transcriptomic data for these tissues were utilized to not only contextualize the model, but also to analyze the nature of the bottleneck genes identified from the model.

From the clustering analysis results of the transcriptome data of the bottleneck genes, it can be concluded that phosphorus-containing pathways and energy-related pathways seem to be consistently involved in bottlenecks when the Arabidopsis roots are exposed to high salt stress conditions. The bottleneck genes were found to be associated with the phosphonate and phosphinate metabolism pathway in each of the root tissues. By further analyzing the protein-protein networks of the clusters it was found that a common bottleneck protein among all cell-types is PBL1, which is a receptor-like cytoplasmic kinase responsible for signaling with calcium and inhibiting root growth during stress conditions. Experimental work published by Kiegle *et al*. (2000) validated that calcium is vital for signaling responses in the Arabidopsis root system when exposed to salt stress. Calcium signaling was also found to be cell type-specific in the roots (Kiegle et al., 2000). Other bottleneck proteins found to be associated with all four root tissue types include ALA2, PECT1, and CCT1, all of which are associated with phospholipid biosynthesis.

To better understand the flux range variation results, it was necessary to consult previous experimental findings. Several studies have shown that during the initial stages of root exposure to salt conditions, the roots go into a "shocked" state where growth and metabolism is halted. However, after several hours of exposure the roots return to a somewhat normal rate of growth (Iyer-Pascuzzi et al., 2011; Machado & Herrgård, 2014; Pirona et al., 2023). From the flux variability results in Figure 7 this is observed especially in the cortex, where all pathways have shrunken flux spaces after 1 hour of salt exposure but have widened flux spaces after 48 hours of salt exposure. However, in the pentose phosphate pathway the formation of D-ribulose 5-phosphate and D-glyceraldehyde 3-phosphate remains suppressed in the cortex under high salt growth conditions. The fatty acid, galactose and purine pathways undergo the same reprogramming as the glycolysis pathway within the cortex.

The endodermis seems to undergo the opposite reprogramming as compared to the cortex. After 1 hour of salt exposure, the majority of the pathways and their associated reactions experience an increase in flux and a widened flux space and a shrunken flux space after 48 hours of exposure, with the exception of the pentose phosphate pathway. It was also observed by Kiegle *et al*. (2000) that the endodermis has a unique response to salt exposure, with higher levels of cytoplasmic free calcium for stress signaling (Kiegle et al., 2000). The endodermis is also responsible for transporting water from the outer cortex and epidermis to the stele, which transports the water, via the xylem, to the rest of the plant (Brady et al., 2007; Dinneny et al., 2008). A separate study showed that the endodermis enhances the deposition of suberin in response to long-term exposure to salt stress, likely to limit water loss from the root (Barberon et al., 2016). Reduced metabolic activity in the endodermis after 48 hours of salt exposure could be due to the suppression of water uptake from the growth media.

The increased flux range in the endodermis experienced by the pentose phosphate pathway is due to an upregulation in the formation of D-ribulose 5-phosphate. A study published by Huang *et al*. (2020) confirmed that D-ribulose 5-phosphate forms an essential part of the cell wall biosynthesis pathway and seems to be upregulated in both the endodermis and epidermis during growth in high salt conditions (Huang et al., 2020). The upregulation of cell wall biosynthesis would allow for cells to maintain internal water retention during high salt growth conditions and provide better support for the root structure (Iyer-Pascuzzi et al., 2011; Kiegle et al., 2000; Zabotina et al., 2012).

Unlike the other cell types, the epidermis was found to have similar metabolic behavior after both 1 hour and 48 hours of high salt growth conditions. All pathways, except the pentose phosphate pathway, were found to have reduced flux spaces in response to salt conditions. However, like the endodermis, the formation of D-ribulose 5-phosphate was upregulated in the pentose phosphate pathway after both 1 hour and 48 hours of salt exposure. The study published by Dinneny *et al*. (2008) confirms that the epidermis is the only cell type in the roots for which gene expression is highly responsive to environmental conditions (Dinneny et al., 2008). The metabolic reprogramming by the stele cells under high salt conditions is very similar to that of the endodermis. Considering that the stele contains both the xylem and the phloem, which are responsible for transporting water and nutrients to the rest of the plant, it can be expected that metabolic activity will be suppressed under prolonged exposure to high salt conditions and result in reduced flux spaces in all pathways after 48 hours of salt exposure.

Considering the overall impact of high salt growth conditions, the different tissue types of the roots undergo unique metabolic reprogramming to ensure that the plant survives. The cortex undergoes an immediate metabolic suppression upon initial exposure to salt conditions, however it is the tissue most likely responsible for root growth, as evidenced by increased synthesis of sugar and organic compounds after prolonged exposure to salt conditions. The metabolic activity in the endodermis, epidermis, and stele is suppressed after prolonged salt exposure, to possibly provide a counter-pressure to the external salt conditions while maintaining internal functionality. The pentose phosphate pathway in the endodermis and epidermis immediately starts producing more D-ribulose 5-phosphate that aids in cell wall biosynthesis to allow for thicker cell walls, which may lead to better water retention in the cells and thickened layers of protection for the roots themselves. Comparing the flux range variation results of AraRoot to previous experimental findings, these results show that AraRoot can accurately capture the metabolic reprogramming that occurs in the root tissues during growth in high salt concentrations.

## 5 Conclusion

The development of the root tissue-specific GEM, AraRoot, highlights the importance of GEMs in studying the metabolic behavior of plants. Not only does the model capture comprehensive biomass formation of the Arabidopsis root system, but it also succeeds in providing valuable insights into the metabolic reprogramming that occurs in different root tissues during exposure to high salt concentrations. Our work highlights the importance of the cortex to root growth in stress conditions, while also capturing the role of the endodermis and epidermis in producing thicker cell walls to allow for better water retention AraRoot could be used to further predict the metabolic responses of the root system to other stress conditions, such as temperature, pH, drought, and nutrient starvation. The methodology proposed here could also be used to develop GEMs for the root systems of major crops. This would allow for further insights into how crops could be improved in terms of resilience to environmental stress and yield.

## Supporting information

Supplemental File 1: AraRoot biomass composition.

Supplemental File 2: Mathematical formulation of all algorithms utilized during model reconstruction and analysis.

Supplemental File 3: Protein-protein interaction networks of the bottleneck genes in each root tissue.

Supplemental File 4: Compilation of the custom database for adding missing reactions, including genes found in roots of soybean, rice, and tomato and their associate Arabidopsis homolog genes.

## Acknowledgements

The authors would like to thank Professor Olga Zabotina from Iowa State University for the insightful discussions regarding hemicellulose formation. We would also like to thank our colleagues in the SSBio laboratory in Lincoln for their insights into reconstructing GEMs.

## Contributions by the Authors

L.E.: Conceptualized problem, reconstructed model, performed analyses, analyzed results, and wrote the original draft. N.A.: Performed analyses, aided in analyzing results, aided in writing of manuscript. M.D.Y.-N., B.J.N., E.E.S.: Provided insights into the metabolism of Arabidopsis, reviewed and edited manuscript R.S.: Procured funding, supervised, provided insights into the methodology, reviewed and edited manuscript.

