## Supplemental File 2 for "AraRoot - A Comprehensive Genome-Scale Metabolic Model for the Arabidopsis Root System"

**FLUX BALANCE ANALYSIS**

Flux Balance Analysis (FBA) is a mathematical approach used to analyze the flow of metabolites through a metabolic network. It is commonly applied in systems biology to predict the growth rate of an organism and the production rates of metabolites. FBA is formulated as a linear programming (LP) problem maximizing or minimizing an objective function subjected to a number of constraints. The most common objective function used in Flux Balance Analysis (FBA) of metabolic networks is the maximization of the biomass reaction flux.

*maximize z = v_biomass_	[FBA]*

*subject to*

$$\sum_{j\in J} S_{ij}v_{j}=0, \forall i \in I$$

$${LB}_{j}\leq v_{j}\leq{UB}_{j}, \forall j \in J$$

*V_j_* $\mathbb{\in R}$

Where,

z: The objective function, which is the biomass reaction flux in this context. The goal of the FBA is to maximize this value, representing the growth rate of the organism.

$S_{ij}$: The stoichiometric coefficient of metabolite *i* in reaction *j*. The stoichiometric matrix *S* is a matrix where rows correspond to metabolites and columns correspond to reactions.

$v_{i}:$ The flux through reaction *j*. It represents the rate at which the reaction occurs.

${LB}_{j}$: The lower bound of the flux for reaction *j*. It sets the minimum rate at which reaction *j* can occur.

${UB}_{j}$The upper bound of the flux for reaction *j*. It sets the maximum rate at which reaction *j* can occur.

**FLUX VARIABILITY ANALYSIS**

Flux variability Analysis (FVA) determines the range of possible flux values for each reaction in a metabolic network while still achieving a given objective value. The flux of each reaction in the network is maximized and minimized, one at a time, while fixing the biomass flux at some fraction f of the optimal value obtained from FBA ($v_{biomass}^{max}$).

*maximize (and minimize) z = v_j_	[FVA]*

*Subject to*

$$\sum_{j\in J} S_{ij}v_{i}=0, \forall i \in I$$

$${LB}_{j}\leq v_{j}\leq{UB}_{j}, \forall j \in J$$

*v_biomass =_ f* $v_{biomass}^{max}$

*v_j_* $\mathbb{\in R}$

**SHADOW PRICE**

In the context of FBA, the shadow price represents the sensitivity of the objective function to changes in the availability of a particular metabolite. It is the dual variable associated with the mass balance constraint for that metabolite. We consider the dual problem of our primal solution.

Primal

*maximize z = v_biomass_	[FBA]*

*subject to*

$$\sum_{j\in J} S_{ij}v_{j}=0, \forall i \in I$$

$${LB}_{j}\leq v_{j}\leq{UB}_{j}, \forall j \in J$$

*V_j_* $\mathbb{\in R}$

Dual

$$minimize z= \sum_{i\in I} 0\lambda_{i}+ \sum_{j\in J} \left( -LB_{j} \right)\mu_{j}^{LB}+ \sum_{j\in J} \left( -UB_{j} \right)\mu_{j}^{UB}$$

*Subject to*

$$\sum_{j\in J} S_{i}\lambda_{i}+ \mu_{j}^{UB}-\mu_{j}^{LB} =0 \forall j\in J-(biomass)$$

$$\sum_{j\in J} S_{ibiomass}\lambda_{i}+ \mu_{biomass}^{UB}-\mu_{biomass}^{LB} =1$$

$$\mu_{j}^{UB},\mu_{j}^{LB}\geq0, \forall j\in J, \lambda_{i}\mathbb{\in R, \forall}i\in I$$

Where $,\lambda_{i}$ is the shadow price for the metabolite *i*

**THERMODYNAMICALLY INFEASIBLE CYCLES**

Thermodynamically Infeasible Cycles (TICs) are sets of reactions that, when taken together, result in a cycle that violates the second law of thermodynamics. This implies that the net flux through the cycle would allow perpetual motion, which is not physically feasible. FBA calculations are limited solely by stoichiometry, which can lead to an unlimited flux through TICs without any external energy input, effectively creating a second-kind perpetual motion machine, which is thermodynamically impossible. The identification and elimination of TICS is important to prevent unbounded metabolic flows and thermodynamically infeasible flux distributions.

*maximize (and minimize) z = v_j_	[TIC]*

*Subject to*

$$\sum_{j\in J} S_{ij}v_{j}=0, \forall i \in I$$

$${LB}_{j}\leq v_{j}\leq{UB}_{j}, \forall i \in I$$

*v_jex_ = 0*

If any flux *v_j_* has a value other than zero, the corresponding reaction j is considered as part of a TIC.
