## Supplemental File 3 for "AraRoot - A Comprehensive Genome-Scale Metabolic Model for the Arabidopsis Root System"

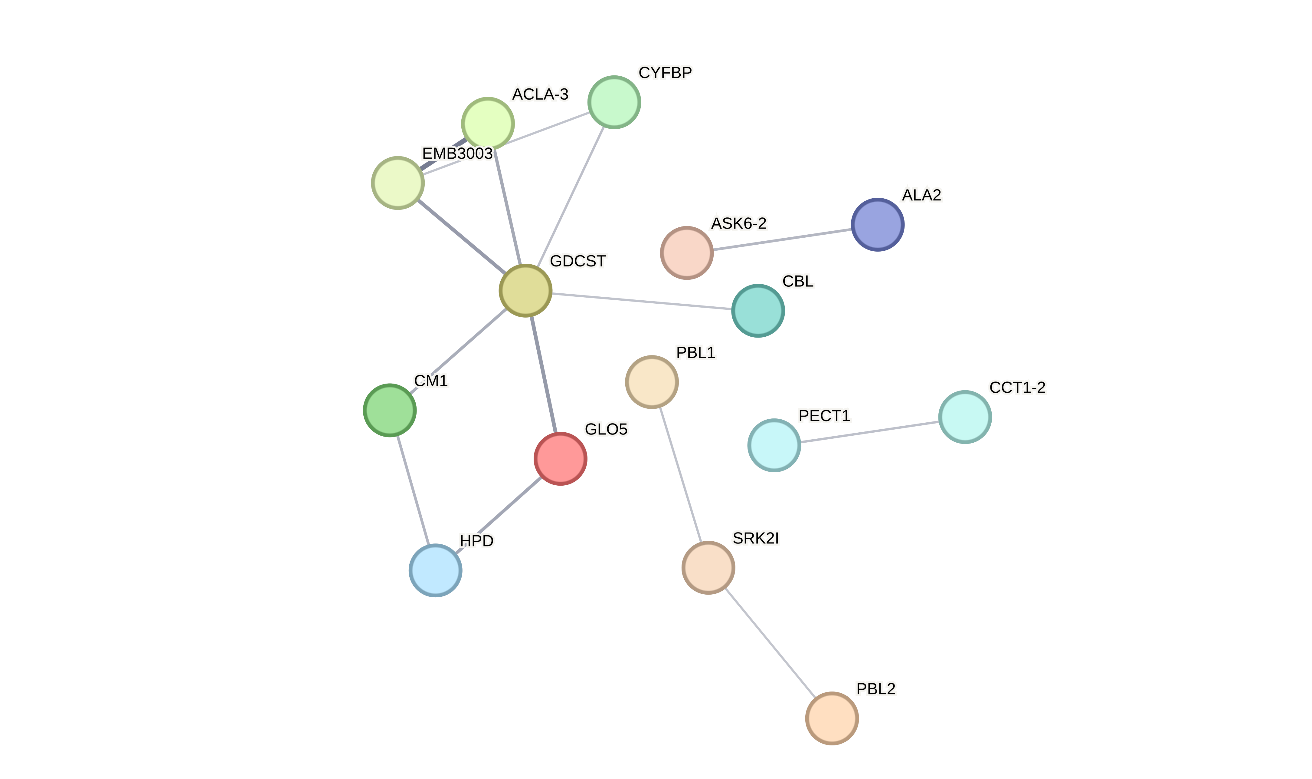

Figure S1: Protein-protein interaction of bottleneck genes in Cortex.

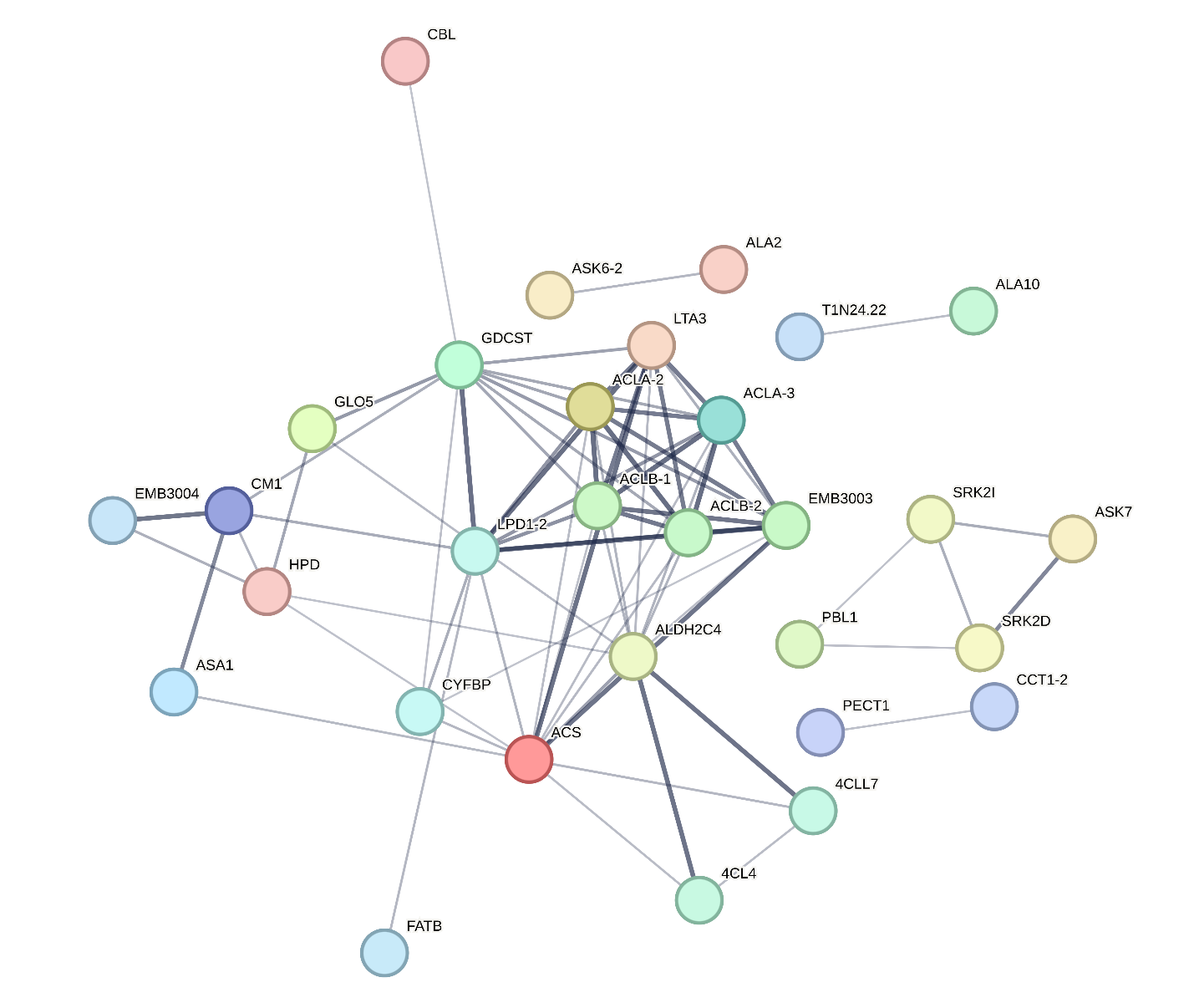

Figure S2: Protein-protein interaction of bottleneck genes in Endodermis.

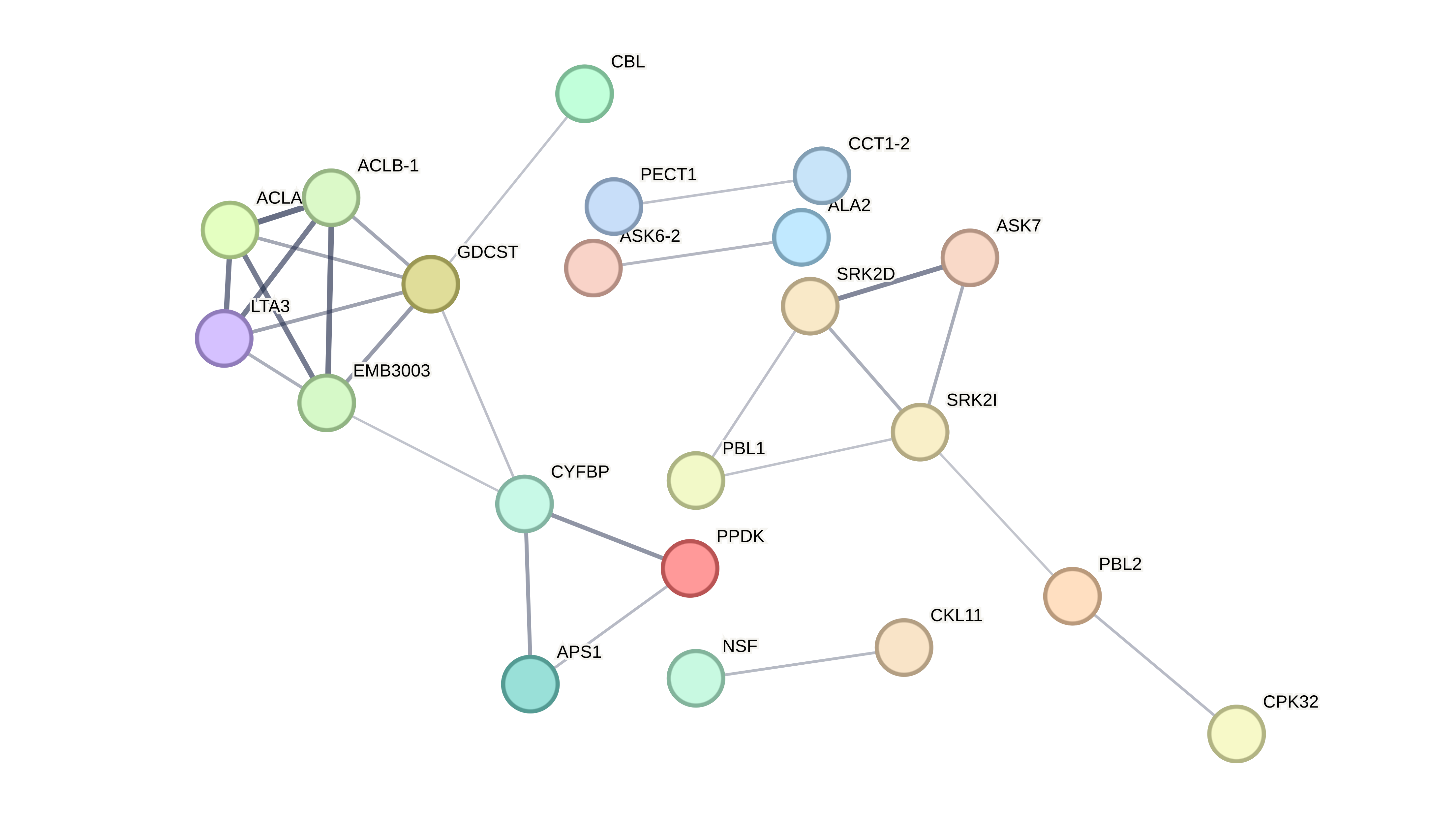

Figure S3: Protein-protein interaction of bottleneck genes in Epidermis.

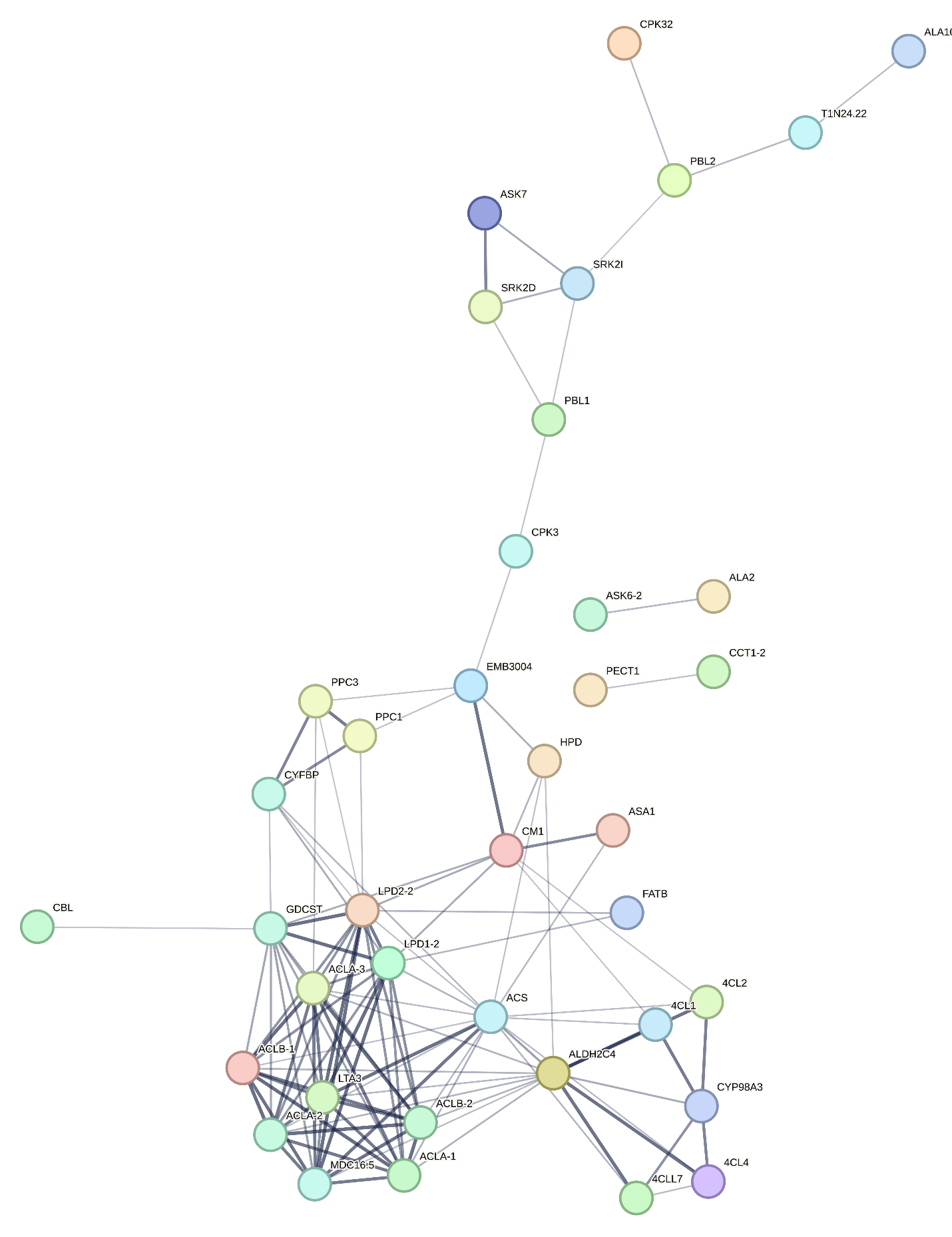

Figure S4: Protein-protein interaction of bottleneck genes in Stele.
